# Efficacy of Distributed Training Depends on the Dorso-Lateral Striatum

**DOI:** 10.1101/2021.03.17.435609

**Authors:** Valentina Mastrorilli, Eleonora Centofante, Federica Antonelli, Arianna Rinaldi, Andrea Mele

## Abstract

Distributed training has long been known to lead to more robust memory formation as compared to massed training. Here we demonstrate that distributed and massed training differentially engage the dorsolateral and dorsomedial striatum and optogenetic priming of dorsolateral striatum can artificially increase the robustness of massed training to the level of distributed training, identifying a novel therapeutic avenue for memory enhancement.

---

Practice is fundamental to learning, and increasing practice is essential to transfer learning into more stable memories. Interestingly, memory improves not only with the increase in the number of repetitions, but also when repetitions are spaced in time. This phenomenon, termed *distributed practice effect*, has long been known by psychologists and has been widely studied in both basic and applied research due to its relevance for education, therapy, and advertising^1,2^. However, its neurobiological underpinnings are still poorly understood. Optimization of memory based on increased number of training trials has been shown to depend on the serial recruitment of functionally distinct neuronal circuits. This is well established for skill and goal directed behavior in the striatal complex. For example, the dorso-medial striatum (DMS) is engaged during the initial phases of skill learning and in the rapid acquisition of action-outcome contingencies^3,4^. In contrast, the dorsolateral striatum (DLS), is critical at later stages of acquisition when performance reaches its asymptotic form^3,4^. Although the striatal complex has been traditionally less investigated in the context of spatial memory^5–7^ a similar serial engagement of the two striatal subregions has also been reported with increasing spatial training in the Morris water maze, in both humans and rodents^7^. Thus, a convergence of empirical evidence supports the view that, regardless of the kind of information to be acquired, striatal compartment-specific activity might predict formation of more enduring memories acquired through extensive training.

Exploiting the striatal complex as a model system here we explored the notion that increased memory efficiency occurring after distributed training might depend upon recruitment of the DLS, in analogy to what has been suggested for prolonged training. To this aim we developed, in mice, a distributed training protocol in the spatial version of the Morris water maze (sMWM). Considering the suggestion that irregular intervals might be more effective in optimizing memory^8^, we selected a three days training procedure consisting in 2 sessions per day, over 3 consecutive days, with a 4h within day interval. To test the efficacy of the distributed training protocol on the acquisition and storage of spatial information, we compared distributed training groups with groups trained with a massed protocol, consisting in the same number of sessions administered consecutively with an inter-session interval of 10 to 15 min^9,10^. Separate groups of mice, trained with the two protocols, in the sMWM, were tested at two different time intervals (24 h and 14 days) after the last training session (Fig. 1a). Probe trial 24 h after training did not reveal any difference between the two groups, both being able to correctly reach the platform location (Fig. 1b; Extended Data Fig. 1a). On the contrary, 14 d after training mice of the massed trained group spent a similar amount of time exploring the four quadrants, demonstrating that spatial memory declined over time. Mice trained with the distributed protocol, instead, maintained intact memory for the correct quadrant location (Fig. 1c; Extended Data Fig. 1b). This difference cannot be attributed to an effect of the training protocol on the learning capabilities, since no differences were found in the latencies to locate the platform during training (Extended Data Fig. 2a,b) or in the progression of the navigational strategies deployed^11^ (Extended Data Fig. 2c; Extended Data Table1) by the two groups across sessions. These findings demonstrate the efficacy of distributed training in creating durable spatial memories, replicating the increased memory performance observed in humans when inter-repetitions lag increases^2^.

**Figure 1.**
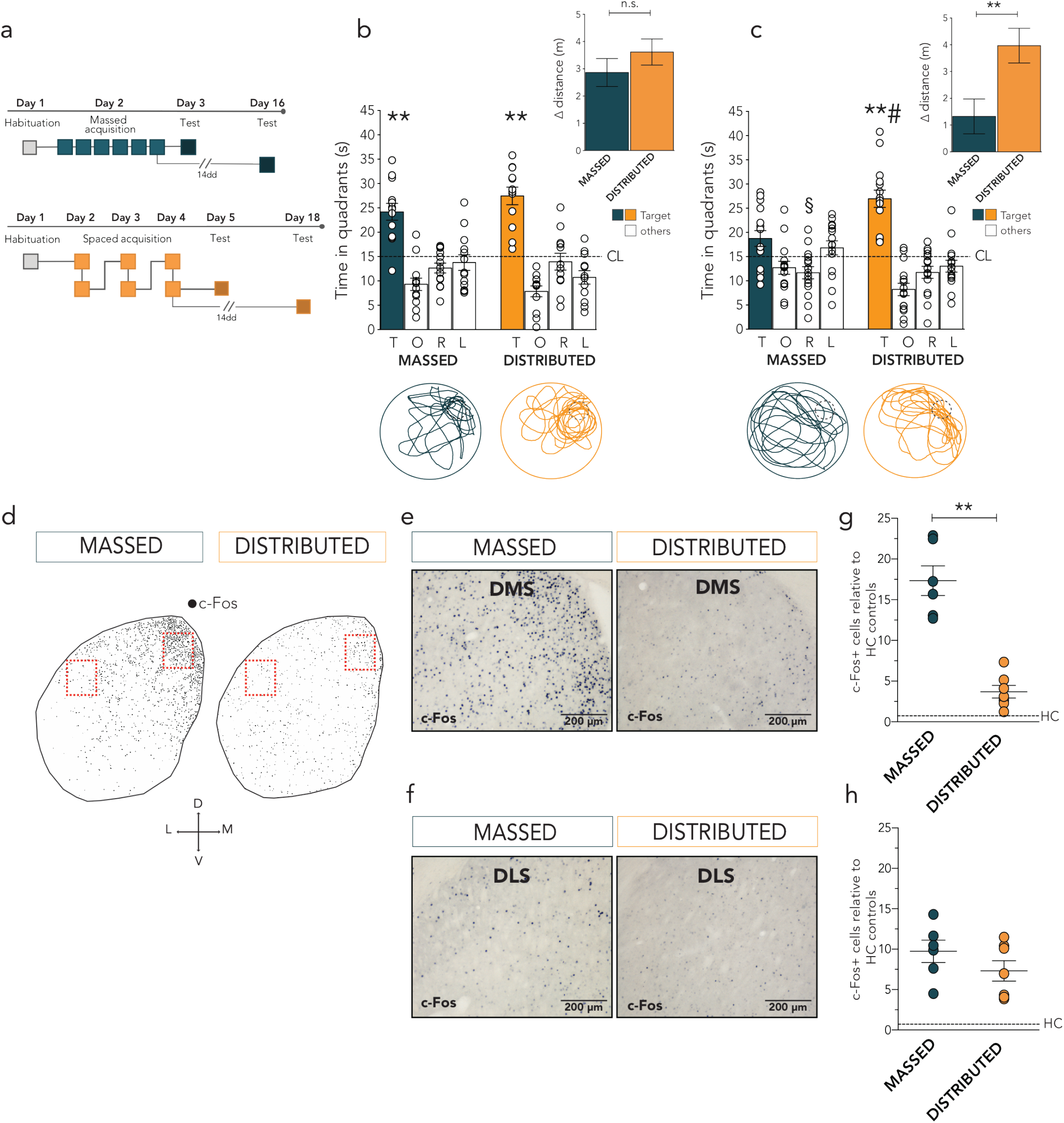
Distributed training increased memory stability and c-Fos expression in the DLS. **a**, Schematic of the experimental design. **b**, Mice trained with the massed (n=13) or the distributed (n=12) protocol were equally able to locate the target quadrant 24h after the last training session (two-way ANOVA repeated measure: quadrant preference F_(3, 69)_=38.41, p<0.0001; protocol F_(1, 23)_=1.53, p=0.228; quadrant preference x protocol F_(3, 69)_=1.35, p=0.265). **c,** 14 days after last training session mice trained with the distributed (n=15), but not the massed (n=15) protocol maintained intact ability to locate the target quadrant (two-way ANOVA repeated measure: quadrant preference F_(3, 84)_=23.27, p<0.0001; protocol F_(1, 28)_=0.086 p=0.770; quadrant preference x protocol F_(3, 84)_=6.374, p=0.0006). Dotted lines represent chance level (CL). The inserts show the distance travelled on test trial expressed as difference (Δ) between target and mean of non-target quadrants, for each group (t_23_=1.063, p=0.298, 24 h and t_28_=2.883, p=0.0075, 14 days after last training session; unpaired *t*-test). Bottom panels: representative path from massed and distributed trained animals. *p<0.05 target vs right, opposite, left; §p<0.05 vs target (within groups, Tukey HSD), # p< 0.05 target vs target (between groups, Tukey HSD). **d,** Mapping of c-Fos immunoreactivity in the whole dorsal striatum of two representative mice trained with the two protocols. **e, f,** Representative images showing training-induced c-Fos immunoreactivity in the DMS **(e)** and in the DLS **(f)** 1 h after massed or distributed training in the sMWM. Scale bar: 200 μm. **g, h,** Quantification of c-Fos expression in DMS and DLS relative to holding cage (HC) mice (n=12). **g,** Cell activity in the DMS was significantly increased in the massed (n=6) compared to distributed (n=7) trained group (U=0.0, p=0.0012). **h,** In the DLS trained mice showed higher level of c-Fos expression compared to HC controls, no significant differences were found between mice trained with the two protocols (U=11, p=0.18). Scatter plots represent mean ± SEM. ** p<0.01 (Mann-Whitney).

To provide direct evidence supporting the hypothesis that the different dorsal striatal domains might be differentially engaged by massed or distributed spatial training, we explored learning-induced changes in neuronal activity, by means of c-Fos labeling, in the DMS and DLS after training with the two protocols in the sMWM (Extended Data Fig. 3). To verify the specificity of the effects, further groups of mice were trained in the cue version of the MWM (cMWM) (Extended Data Fig. 3). A group of home caged (HC) controls was used to normalize data obtained from spatial trained mice. Mirroring what has been reported on the contribution of the DMS and DLS after short or prolonged training^7^, we found that compared to HC controls the DMS was specifically activated by massed but not distributed training (Fig. 1d,e,g; Extended Data Fig. 4). On the contrary, increased c-Fos expression in the DLS was independent of the training protocol (Fig. 1d,f,h; Extended Data Fig. 4). Importantly, with the notable exception of the DMS after distributed training, that showed similar c-Fos expression in the cue and the spatial groups, cell activity-dependent labeling was higher in the spatial trained groups compared to those trained in cue version of the task (Extended Data Fig. 4) independently on the training protocol.

To assess whether increased neuronal activity in DMS after massed training and in the DLS after massed and distributed training had a causal role in memory storage, we performed a loss of function manipulation of the two striatal domains on mice trained with the two protocols. To this aim the AMPA-R antagonist, NBQX, was administered immediately before testing 24h after the last training sessions, a time point at which both massed and distributed training induced effective memory. We first examined the effects of DMS and DLS manipulations on mice trained with the massed protocol (Fig. 2a). As expected, both groups of vehicle injected mice were perfectly able to locate the platform. However, differently to what could be expected from c-Fos labeling data, pre-test administration of NBQX impaired mice ability to locate the platform only when injected in the DMS, but not in the DLS (Fig. 2b,c; Extended Data Fig. 5a-d; 6a,b; 7a) compared to vehicle controls. Notably, the opposite effect was found when testing the effects of DMS and DLS manipulations on spatial memory acquired with the distributed protocol (Fig. 2a). Mice receiving pre-test NBQX in the DMS, in fact, showed intact ability to locate the platform compared to vehicle injected controls (Fig. 2d; Extended Data Fig. 5e,f; 6c). On the contrary, pre-test NBQX administrations in the DLS impaired mice performance on the probe trial 24h after the last training session (Fig. 2e; Extended Data Fig. 5g,h; 6d; 7b). Overall, these data demonstrate that the two striatal domains are both involved in the processing of spatial information, providing a causal role to neural activation reported in the previous experiment as well as in imaging studies in humans and rodents^7^, but more important they suggest a double dissociation in their involvement depending on the timing rule of the learning protocol.

**Figure 2.**
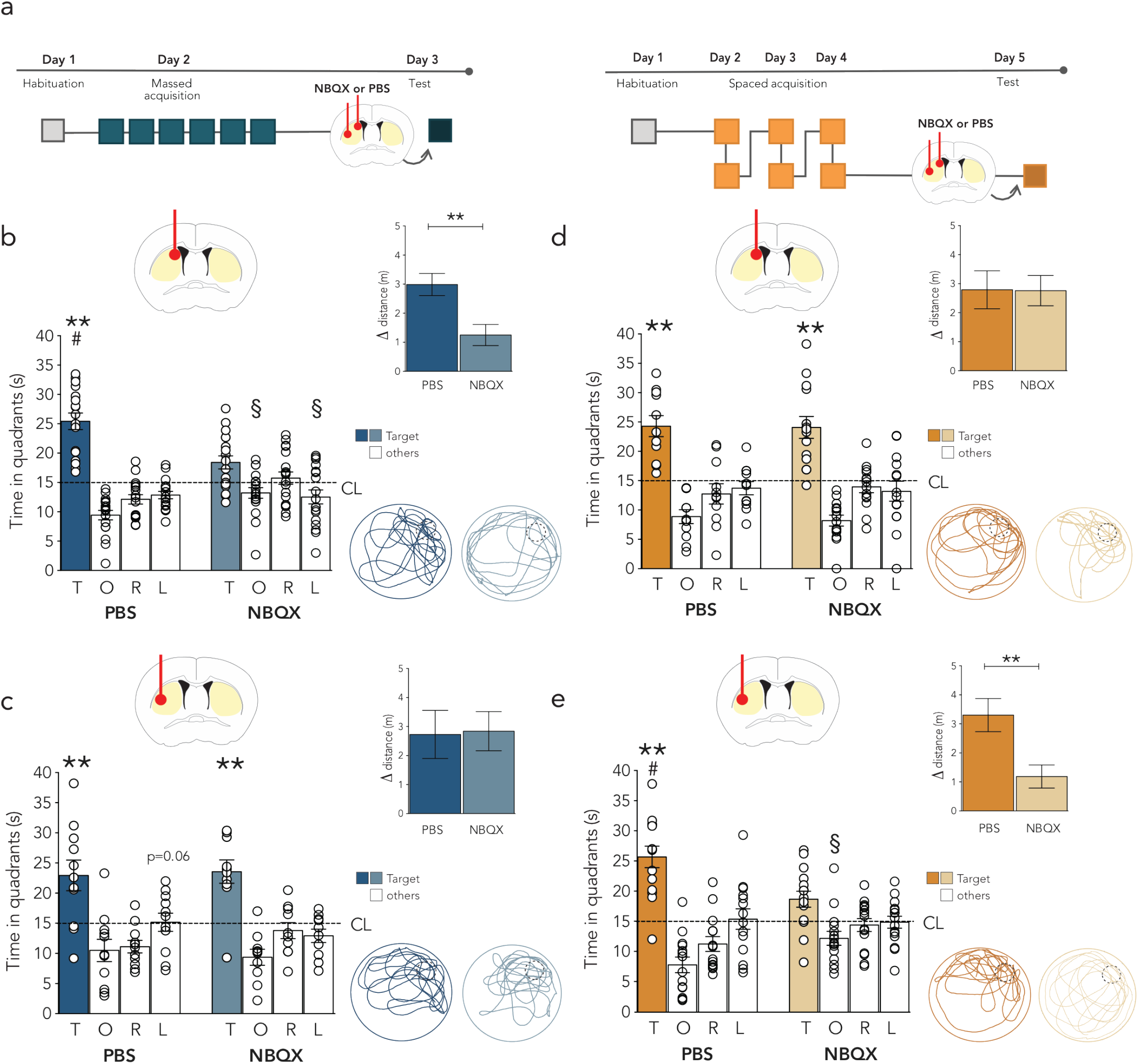
Differential involvement of the DMS and DLS in the retrieval of spatial memory acquired through massed or distributed training. **a,** Schematic of experimental design. **b,** Pre-test administrations of NBQX in the DMS (n=18) impaired mice ability to correctly locate the platform on probe trial 24 h after massed training, compared to vehicle injected controls (n=17), (two-way ANOVA repeated measure: quadrant preference F_(3,99)_=32.63, p<0.0001; treatment F_(1,33)_=0.78, p=0.38; quadrant preference x treatment F_(3,99)_=9.16, p<0.0001). The insert shows the distance travelled on probe trial expressed as difference (Δ) between target and non-target quadrants, in the two groups (t_33_=3.29, p=0.0024; unpaired *t*-test). **c,** Pre-test administrations of NBQX in the DLS (n=10) did not affect mice ability to correctly locate the target quadrant on probe trial 24 h after massed training compared to vehicle controls (n=11) (two-way ANOVA repeated measure: quadrant preference F_(3,57)_=18.15, p<0.0001; treatment F_(1,19)_= 0.41, p=0.53; quadrant preference x treatment F_(3,57)_=0.62, p=0.61). The insert shows the distance travelled on probe trial expressed as difference (Δ) between target and non-target quadrants, in the two groups (t_19_=0.10, p=0.918; unpaired *t*-test). **d,** Pre-test administrations of NBQX in the DMS (n=14) did not affect mice ability to correctly locate the target quadrant on probe trial 24h after distributed training, compared to their vehicle controls (vehicle, n=11) (two-way ANOVA repeated measure: quadrant preference F_(3,69)_=30.02, p<0.0001; treatment F_(1,23)_=0.59, p=0.45; quadrant preference x treatment F_(3,69)_=0.13, p=0.94). The insert shows the distance travelled on probe trial expressed as difference (Δ) between target and non-target quadrants, in the two groups (t_23_=0.035, p=0.972; unpaired *t*-test). **e,** Pretest administrations of NBQX in the DLS (n=15) impaired mice ability to correctly locate the platform on probe trial performed 24 h after distributed training, compared to controls (n=14), (two-way ANOVA repeated measure: quadrant preference F_(3,81)_=22.82, p<0.0001; treatment F_(1,27)_=0.0057, p=0.94; quadrant preference x treatment F_(3,81)_=5.49, p=0.0017). The insert shows the distance travelled on probe trial expressed as difference (Δ) between target and non-target quadrants, in the two groups (t_27_=3, p=0.0057; unpaired *t*-test). Dotted line represents chance level (CL). Representative path from control and treated mice in the different groups are also shown. Histograms represent mean ± SEM. *p<0.05 target vs right, opposite, left; § p<0.05 *vs* target (within groups, Tukey HSD); # p<0.05 (between group, Tukey HSD).

Having shown that interference with DLS neuronal activity impairs retrieval of spatial information acquired through a distributed training, we wondered whether the engagement of this region could be a mechanism responsible for the establishment of more enduring memories. Cell activity dependent labeling of the DLS after massed training suggests that short intertrial intervals provide sufficient stimulation for activation, but not for the instantiation of the memory trace. Based on recent finding demonstrating that increasing the excitability of a subset of neurons increases the probability that those neurons will participate in a memory trace^12^, we reasoned that the artificial priming of cell activity in the DLS could bias the information acquired through massed training within this striatal domain and in this way create a more stable trace. To verify this possibility, we injected in the DLS an AAV carrying a light activated excitatory channelrhodopsin, ChR2(C128S/D156A) that responds to light delivery with a long-lasting effect (29min)^13^ (Fig. 3a). As expected, control mice not receiving light delivery were not able to locate the correct quadrant on probe trial 14 days after massed training in the sMWM (Fig. 3b). On the contrary, light (473Hz) stimulated mice showed intact ability to locate the platform on the remote probe trial despite the massed training (Fig. 3b; Extended Data Fig. 8 a-c; 9). Further control experiments demonstrated the efficacy of the light stimulation in promoting increased neuronal activity in the DLS (Fig. 3c) and ruled out possible effects of light delivery alone on learning (Fig. 3d; Extended Data Fig. 8 d-f; 10). These findings demonstrate that artificial stimulation of the DLS in massed training conditions enable the constitution of a more stable spatial memory representation, artificially mimicking the increased memory efficacy induced by distributed training.

**Figure 3.**
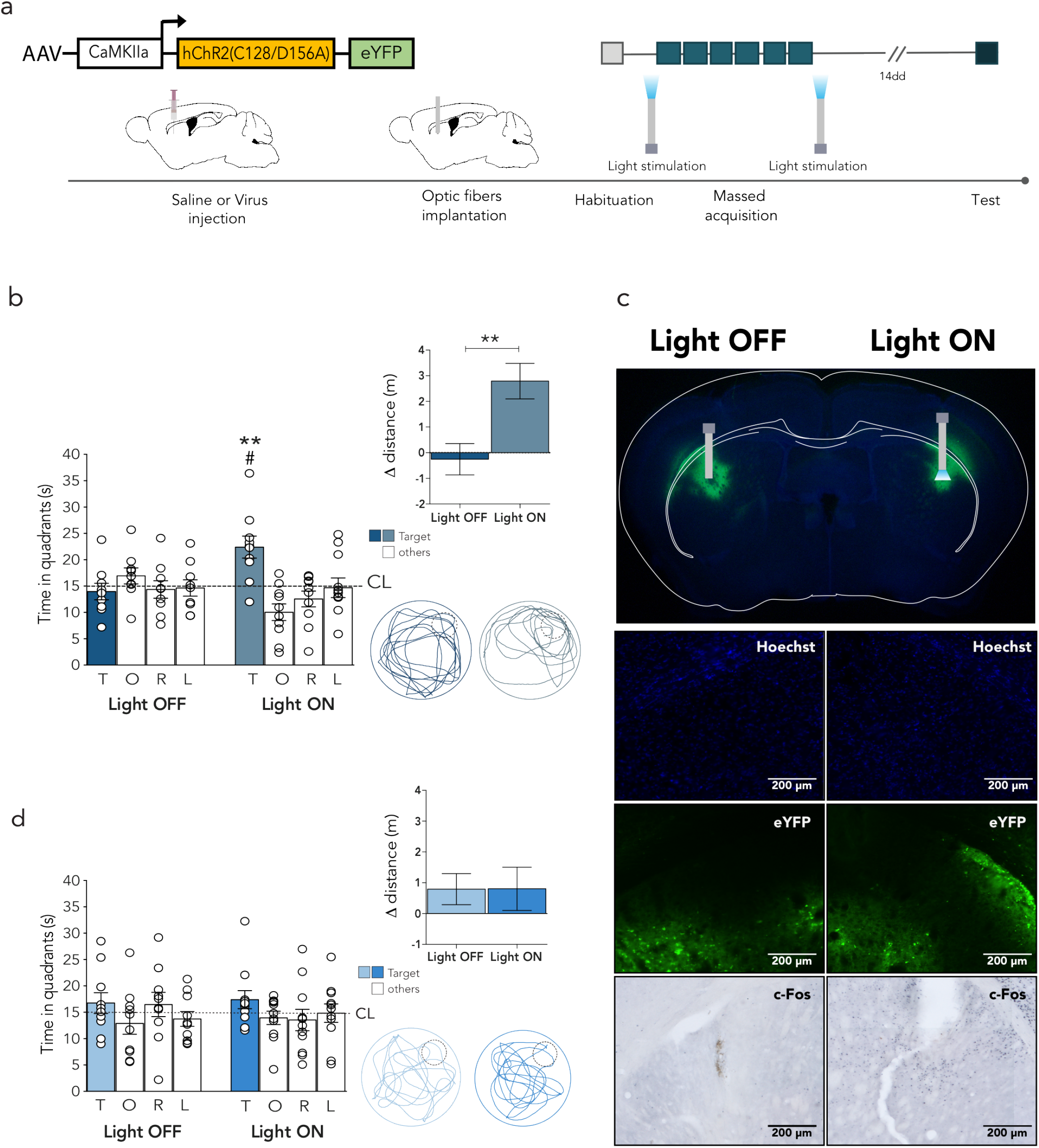
Optogenetic stimulation of DLS enhance memory stability in massed-trained mice. **a,** Schematic illustration of the experimental design. **b**, Light delivery in DLS improved the ability to correctly locate the target quadrant on probe trial 14d after the massed training, in stimulated (n=8) but not unstimulated infected mice (n=9), (two-way ANOVA repeated measure: quadrant preference F_(3,45)_=4.23, p=0.010; treatment F_(1,15)_=1.359, p=0.26; quadrant preference x treatment F_(3,45)_=7.88, p=0.0002). Dotted line represents chance level (CL). The insert shows the mean difference (Δ) between the distance travelled in the target compared to non-target quadrants (t_15_=3.312, p=0.0047; unpaired *t*-test). **c,** Microphotographs showing ChR2-eYFP expression counterstained with Hoechst and representative fiber location in a mouse receiving light delivery unilaterally in the DLS. Representative images of Hoechst, eYFP and c-Fos immunoreactivity in the two emisphere showing increased c-Fos labeling only in the stimulated side. Scale bar: 200 μm. **d,** Light delivery in mice bilaterally administered with saline did not affect performance in the probe trial 14 d after massed training, both groups (light on n=10; light off n=11) being equally unable to locate the target platform 14d after the last training session (two-way ANOVA repeated measure: quadrant preference F_(3,57)_=1.094, p=0.359; protocol F_(1,19)_=0.564, p=0.461; quadrant preference x protocol F_(3,57)_=0.421, p=0.738). Dotted line represents chance level (CL). The insert shows the mean difference (Δ) between the distance travelled in the target compared to non-target quadrants (t_19_=0.0133, p=0.989; unpaired *t*-test). Representative path from control and treated mice in the different groups are also shown. Histograms represent mean ± SEM. *p<0.05 target vs right, opposite, left (within group, Tukey HSD); # p<0.05 (between group interaction, Tukey HSD).

Overall, our findings support our claim that the activation of the DLS after distributed learning has a causal role in the formation of more durable memories. What characterizes distributed compared to massed training is the duration of the off-line resting periods between sessions. By showing that the acquisition of enduring memories through distributed learning requires the engagement of the DLS, our data imply that the temporal dynamics shaping plasticity in this neuronal system might be a key element to understand its ability to generate stable memory representations occurring after both prolonged and distributed training. Moreover, demonstrating that both striatal domains are involved in spatial information processing, but their participation changes based on the temporal spacing of the training sessions, they support the view that differences among memory systems could be addressed based on neurobiological determinants of the processing operations involved, rather than by the kind of information processed^14^.

## Acknowledgements

The authors would like thank C.G. Gross and M.G. Leggio for critically reading the manuscript and for the stimulating discussions, Francesco Grassi, Domenico Cicogna, Claudia Ferrante, Samyutha Rajendran, Mireille Mohamad, for their help with the experiments. This study was supported by a NARSAD independent investigator grant from the Brain and Behavioral Research Foundation (to A.M.) and by grants from the University of Rome “La Sapienza” (to A.M.; A.R.).

## Author contributions

V.M. carried out, designed and analyzed the Fos and the pharmacological experiments; optogenetics experiments were designed and carried out by F.A. and E.C.. The project was conceived by V.M., A.R. and A.M.. The manuscript was written by V.M., A.R. and A.M. with inputs from E.C. and F.A.

## Supplemental information

### METHODS

#### Subjects

The experiments were conducted on naïve CD1 male mice (Charles River, Italy). Mice were at least 7 weeks old, weighting about 35g at the onset of the experiments. Animals were always housed in groups of three-five mice in standard cages (26.8 × 21.5 × 14.1 cm), with water and food *ad libitum*, under a 12 h light/dark cycle and constant temperature (22 ± 1 °C). Behavioral training and testing were conducted during the light period (from 9:00 am to 5:00 pm). All animals were treated in respect to current Italian and European laws for animal care, and the maximum effort was made to minimize animal suffering.

#### Stereotaxic surgery and viral injections

Each mouse was deeply anesthetized with 3% isoflurane (Isovet; Piramal Healthcare, Italy) and secured on the stereotaxic apparatus (David Kopf Instruments, USA) with front teeth incisor bar and lateral zygomatic cups, and eyes were quickly protected with Lacrigel eye gel (Bracco, Italy). For the pharmacological experiment, two stainless-steel guide cannulae (0.50/0.25 × 7 mm; Unimed, Switzerland) were implanted. Cannulae implantation was performed through craniotomies on the skull at the following coordinates relative to bregma, according to the mouse brain atlas^1^: anteroposterior (AP)= +0.30 mm, mediolateral (ML)= ±1.6 mm, dorso-ventral (DV)= −1.3 mm for the dorsomedial striatum (DMS); anteroposterior (AP)= +0.30 mm, mediolateral (ML)= ±2.8 mm, dorso-ventral (DV)= −1.3 mm for the dorsolateral striatum (DLS). Guide cannulas were fixed with acrylic cement (Riccardo Ilic, Italy) to be stably held on the calvarium. After surgery, mice were allowed to recover in their home cages for at least 7 days after surgery.

For the optogenetic experiment, Bi-stable Step Function Opsin (SSFO) AAV5-CamKII-hChR2-eYFP (Gene vector and virus core, Stanford University, USA) was bilaterally inoculated in the DLS at the same AP and ML coordinates reported above, DV= −3.0 mm from the dura. The volume of injection was 0.20 μl per side, delivered at the rate of 0.1 μl/min with the use of glass micropipette. The pipette was lowered to the target site and left in place for additional 5 min to allow diffusion before it was withdrawn. Control animals were injected with the same volume of saline solution (NaCl 0.9%) by using the same procedure. Animals were left to recover for 2 weeks to allow the virus expression. After the recovery period, mice underwent bilateral fibers implantation in the DLS. A custom-made optic fiber (200 nm core diameter; 0.39NA, Thorlabs) was lowered above the injection site (DLS: AP= +0.30 mm, ML=±2.8 mm, DV=-2.5 mm) and secured to the skull with dental cement. Animals were allowed to recover for further 7-9 days before behavioral experiments. All mice were given 2 mg/kg paracetamol as analgesic during the recovery. All injection sites were verified histologically. We only included in the analysis mice with correct placements and minimum 70% of virus expression, limited to the targeted regions.

#### Drug and in vivo focal injection procedure

Drug infusion was performed 20 min before the test phase. A total of 0.25 μl of 0.95 μg/μl of the AMPA receptor antagonist 1,2,3,4-Tetrahydro-6-nitro-2,3-dioxo-benzo[f]quinoxaline-7-sulfonamide disodium salt hydrate (NBQX; Sigma-Aldrich, Italy) was bilaterally infused in the brain at the injection rate of 0.125 μl/min for 2 min. NBQX was dissolved in phosphate-buffered saline (PBS 0.1 M). Control mice were injected with the same volume of PBS. During the injection, the needle (length, 9 mm; diameter 0.25 mm; Unimed, Switzerland) was connected by a plastic tube to a 2-μl Hamilton syringe and mice were awake and free to move in the holding cage. After administration, the injector was left in place for 1 additional minute to allow diffusion.

#### Behavioral procedures

The spatial water maze task (sMWM) consisted of three different phases distributed across three consecutive days: familiarization, training and test. During the whole task, the water was made black with non-toxic paint (Giotto, Italy) and water temperature was kept constant at +22°C. Familiarization consisted of one session of three consecutive trials (intertrial interval: 20 s). No cues were attached to the wall and a 10-cm-diameter platform protruded 1.5 cm above the water surface. The session started with the animal standing on the platform for 20 s. Animals were introduced in a non-target quadrant in a pseudo-random order facing the wall. If they did not reach the platform in 60 s, they were gently led to the platform by the experimenter. The massed training consisted of six consecutive sessions (intersession interval: 10-15 min) of three trials (intertrial interval: 30 s)^2,3^. The distributed version of MWM consisted of six sessions distributed over 3 days, with 2 sessions per day (intersession interval: 4 h) of three trials (intertrial interval: 30 s). The procedure was the same as in the familiarization phase except that the platform was submerged 0.5 cm beneath the surface of the water. During this phase, the pool was surrounded by several extra-maze cues objects placed as reference cues onto black curtains. Depending on the experiment, the probe trial was performed 24 h or 14 days after the last training session and consisted of a single trial. The platform was removed, and mice were allowed a 60-s search for the platform starting from the center of the pool. General training measures included latency to reach the platform and path length. Search strategies during training were analyzed using numerical parameters from swim tracking data (adapted from^4^), and the respective predominant search strategy for each trial was objectively classified by a criterion-based custom-made algorithm. Trials were classified as one of the following ordered strategies (tracking criteria in parentheses): thigmotaxis: >35% of swim distance within closer wall zone (10 cm from pool wall) and total latency to the platform < 25s; random search: total latency >25 seconds; scanning: <75% surface coverage and >35% surface coverage <0.7 s.d. distance to the pool center (scanning area) and total latency >25 seconds; chaining: between 35-70% of time within chaining zone (chaining zone: corridor 20cm wide, centered in the center of platform) and total latency >25 seconds; missing-then-reaching: latency to critical area/latency to the platform =0.2-0.5 and total latency <11s (critical area: 30cm-wide area centered in the center of the platform); directed search: >70% of time in goal corridor (rectangular goal corridor 30 cm wide, centered along direct connection between start and platform positions); focal search: <20 s.e.m. body angle, <0.25 s.d. mean distance to present goal; direct swim: >90% distance in the goal corridor. When we used these definitions and the algorithm, only <2,5% of the trials could not be assigned univocally to one strategy. Probe trial performance was quantified as the time spent, distance travelled or annuli crossed in target compared to other three non-target quadrants.

In the cue version of the water maze task (cMWM) all the distal cues were removed and a single proximal cue was present, a green ball hanging 5 cm above the hidden platform^5^ (Ferretti et al., 2015). The position of the platform and the ball changed across sessions to prevent animals from using spatial bias. Behavioral data from training trials were acquired and analyzed using an automated tracking system (ANY-maze, Stoelting).

#### Video analysis and data collection

All the trials during the familiarization, training and test were recorded trough a camera located above the pool and videos were analyzed with the software AnyMaze-Behaviour Tracking System 5.0 (ANY-maze, Stoelting, USA). Several variables were *a posteriori* scored. To study behavior during the training, the variables scored were escape latency (in seconds) to reach the platform, escape distance travelled to reach the platform (in m). For the probe trial we set with AnyMaze an arena corresponding to the poll circumference. The arena was divided in 4 quadrants (target, opposite, left and right) and 4 annuli (target, opposite, left and right), defined as an area of 18 cm diameter surrounding the center of the platform for each quadrant. The following variables were scored during the probe phase: time spent in each quadrant (in seconds), distance travelled in each quadrant (in m), annulus crossing (frequency of each annulus crossing).

#### Immunohistochemistry: c-Fos staining

One hour after completing the last training session in the Morris water maze, each animal was deeply anaesthetized with mixture of Zoletil (500 mg/kg, Virbac Italia) and Xylazine (100 mg/kg, Bayer, Germany) and transcardially perfused with 40 ml of saline solution (NaCl 0.9%) followed by 40 ml of 4% formaldehyde. Brains were rapidly removed and post-fixed for 24 h in 4% formaldehyde in PBS and then transferred to 30% sucrose in PBS until sectioning. 30 μm-coronal sections were obtained using a freezing microtome (Leica CM 1950, Leica Microsystems, Wetzlar, Germany) and were stored at −20 °C in cryoprotectant solution. For c-Fos detection in the optogenetic experiment, the day after the test animals underwent to unilateral optical stimulation, left undisturbed in their holding cages for one hour, then anesthetized and perfused as described above. 40 μm-coronal sections were collected by using a freezing microtome (Leica CM 1950, Wetzlar, Germany) and were stored at −20 °C in cryoprotectant solution.

For both experiments sections to be processed for c-Fos-immunoreactivity (c-Fos-IR) were transferred in 0.1 M phosphate-buffered saline (PBS, pH 7.4) and washed several times. Briefly, free-floating sections were incubated for 5 min in 3% hydrogen peroxide in PBS and rinsed three times in PBS containing 0.1% Triton X-100 (PBST). After 1 h of incubation in PBST containing 1% BSA and 1% NGS (PBST–BSA–NGS) sections were incubated overnight in anti-phospho-c-Fos rabbit monoclonal antibody (5348S; Cell signaling Technology, USA) diluted 1:8000 in PBST–BSA–NGS, at 4 °C and with constant orbital rotation. Sections were washed three times in PBST and incubated in biotinylated secondary antibody diluted 1:500 in PBST-1% BSA (goat anti-rabbit IgG; Vector Laboratories, USA) for 2 h at room temperature. After three washes in PBST sections were transferred for 1 h in avidin–biotinylated peroxidase complex diluted 1:500 in PBST (ABC Kit; Vector Laboratories, USA) and rinsed three times in PBS. Finally, the reaction was visualized using nickel intensified diaminobenzidine (DAB peroxidase substrate kit, Vector Laboratories, USA). The reaction was stopped after exactly 4 min by washing with 0.1 M PBS (pH 7.6). After several rinses in PBS sections were mounted on slides, dehydrated through a graded series of alcohols, cleared and coverslipped for microscopical examination. Sections from groups to be directly compared were processed at the same time and using the same conditions and reagents in order to reduce variability. In all the experiments, the number of cells displaying c-Fos-IR was measured in DMS and DLS (from bregma +0.50 mm to bregma +0.02 mm). Regions were defined according to the mouse brain atlas^1^. At least three to five non-consecutive coronal sections were stained and digitized bilaterally for each brain region, for each subject. Digital images were acquired at 10× magnification, using a microscope (Nikon eclipse 80i) equipped with a CCD camera. After images acquisition, counting of the stained cells was carried out using the public domain software Image J (NIH Image, USA/). Briefly, for each region, stained nuclei were automatically detected based on their intensity of staining relative to background and their size. These parameters were set based on previous extensive comparisons between automatic and manually generated counts. Counts from both hemispheres and from all rostro-caudal levels were averaged per subject, and subsequently per group (naïve, cue and spatial).

#### Digital reconstruction

Following c-Fos staining, one representative sections (+0.38 mm from bregma) from one representative mouse for each training protocol was digitized for the reconstruction of c-Fos expression throughout the whole striatum. Digital images were acquired at × 10 magnification, using a Nikon microscope (Nikon eclipse 80i) equipped with a digital camera (DXM1200F). Single images of 2560 × 1920 pixel were stitched together in a mosaic view with the use of Adobe Photoshop (Adobe Photoshop CC 2019). Images were then processed with a custom made script in MatLab.

#### Optical stimulation

Mice were acclimatised to their holding cages for at least 30 min before the light stimulation. One light pulse of 1 second (pulser Prizmatix, USA) was delivered for both hChR2 and saline-injected mice 10 minutes before the beginning of the training and right after the end of the training in the sMWM. During stimulation, mice were free to move in their holding cages. At the end of the stimulation mice were detached and returned to their waiting cages for 10 and 30 minutes before and after the training respectively. Besram Technology Inc. laser (blue 473 nm, Wuhan Besram Technology Inc, China) with of 0.8-1 mW at the fiber tip (PM100D power sensor, Thorlabs, Germany), was used as light source.

#### Histology and viral diffusion

To verify the location of the injections in the pharmacological experiments and optic fiber location in the optogenetic experiment, at the completion of testing animals were anesthetized with isoflurane followed by cervical dislocation. Brains were removed and post-fixed in 4% formaldheyde (4°C). 90-μm thick coronal slices were cut using a microtome (Leica Microsystem, Germany), collected on gelatin-coated slides and stained with cresyl-violet (Sigma-Aldrich, Italy). They were then analyzed with the use of a stereomicroscope and the most ventral point of the tip of the injector or the optic fiber was identified. Injections or fiber location sites were estimated with reference to a mouse brain atlas. Only mice with correct placement in the target regions were included in the analysis. Illustration of coronal sections from single animals are represented for each experiment (Extended Data Figs. 7, 9, 10).

To verify the viral diffusion and optical fibers placements in the optogenetic experiments, mice were anesthetized with an overdose of mixture of Zoletil (500 mg/kg, Virbac Italia) and Xylazine (100 mg/kg, Bayer, Germany) and transcardially perfused with 0.1 M ice-cold PBS followed by 4% formaldehyde immediately after the test phase. Brains were post-fixed overnight at 4 °C in 4% formaldehyde and then cryopreserved in 30% sucrose. Coronal slices containing DLS (40-μm thick) were cut using a freezing microtome (Leica Microsystem, Germany) and collected in PBS. For virus detection, slices were incubated 5 min with Hoechst (Termofisher, USA) diluited 1:2000, immediately mounted and covered with Fluoromount aqueous mounting medium (Sigma-Aldrich, Italy). Immunofluorescence images were acquired with a fluorescence microscope (Nikon Eclipse TE300) with 2x objective under 488nm laser/eYFP setting. Total virus diffusion was outlined and determined for each section with the use of ImageJ (NIH) and Adobe Illustrator (Adobe Systems Incorporated, USA). Only mice showing AAV diffusion in at least the 70% of total DLS extension and a correct optic fibers placement were included in statistics (Extended Data Figs. 9). To verify viral diffusion in unilateral-stimulated mice, slices consecutive to the ones used for c-Fos staining were used.

#### Data collection and statistical analysis

All data were represented as mean ± SEM. For statistical analysis, Statistica Software (Dell Software, Oklahoma), Prism (GraphPad Software, USA) and SPSS (IBM, Italy) were used. Group differences were considered statistically significant when p< 0.05.

For the analysis of training for all MWM experiments, escape latency and path length data were analyzed using two-way repeated measures ANOVA with procedure (MASSED or DISTRIBUTED), pre-test treatment (NBQX or PBS) or light stimulation (LIGHT ON or LIGHT OFF) as between groups factor and session (six levels: session 1 to 6) as repeated measure. If the interaction between factors was not significant a one-way repeated measure ANOVA with session (six levels: session 1 to 6) as repeated measure was performed on the two groups independently. To compare the navigational strategies in distributed vs massed, a multinomial logistic regression was conducted using 4 navigational strategies (spatial, spatial non specific, non spatial and non classified) predictors to predict the classification on the multinomial dependent variable offence type (massed vs distributed).

For probe trial time, distance and annuli were analyzed by using two-way repeated measure ANOVA with procedure (two level, DISTRIBUTED or MASSED), pre-test treatment (two level, PBS or NBQX) or light stimulation (two levels, LIGHT ON or LIGHT OFF) as between groups factor and quadrants (4 levels: target, opposite, right and left) as repeated measure. Group differences were considered statistically significant when p<0.05. If this was not the case, the repeated measures were analyzed independently in each group (PBS, NBQX or DISTRIBUTED, MASSED or LIGHT ON, LIGHT OFF) through a one-way ANOVA with quadrants (4 levels: target, opposite, right and left) as repeated measure. If this was statistically significant Tukey HSD post-hoc analysis was used. Difference between the target distance and the mean of non-target quadrants was analyzed by using *paired* t-test. To compare c-Fos expression, normalized on HC controls, in mice from the massed and distributed trained groups a Mann-Whitney non-parametric test was used. To compare c-Fos expression (c-Fos/mm^2^) in HC, spatial and cue trained mice with the two protocols one-way ANOVA was used with training (HC, spatial, cue) as between-groups factor. If this was statistically significant Tukey HSD post-hoc analysis was used. Principal Component Analysis (PCA) was run to compare different stainings by using an open-source web tool (https://biit.cs.ut.ee/clustvis).

---

**Extended Data Fig. 1.**
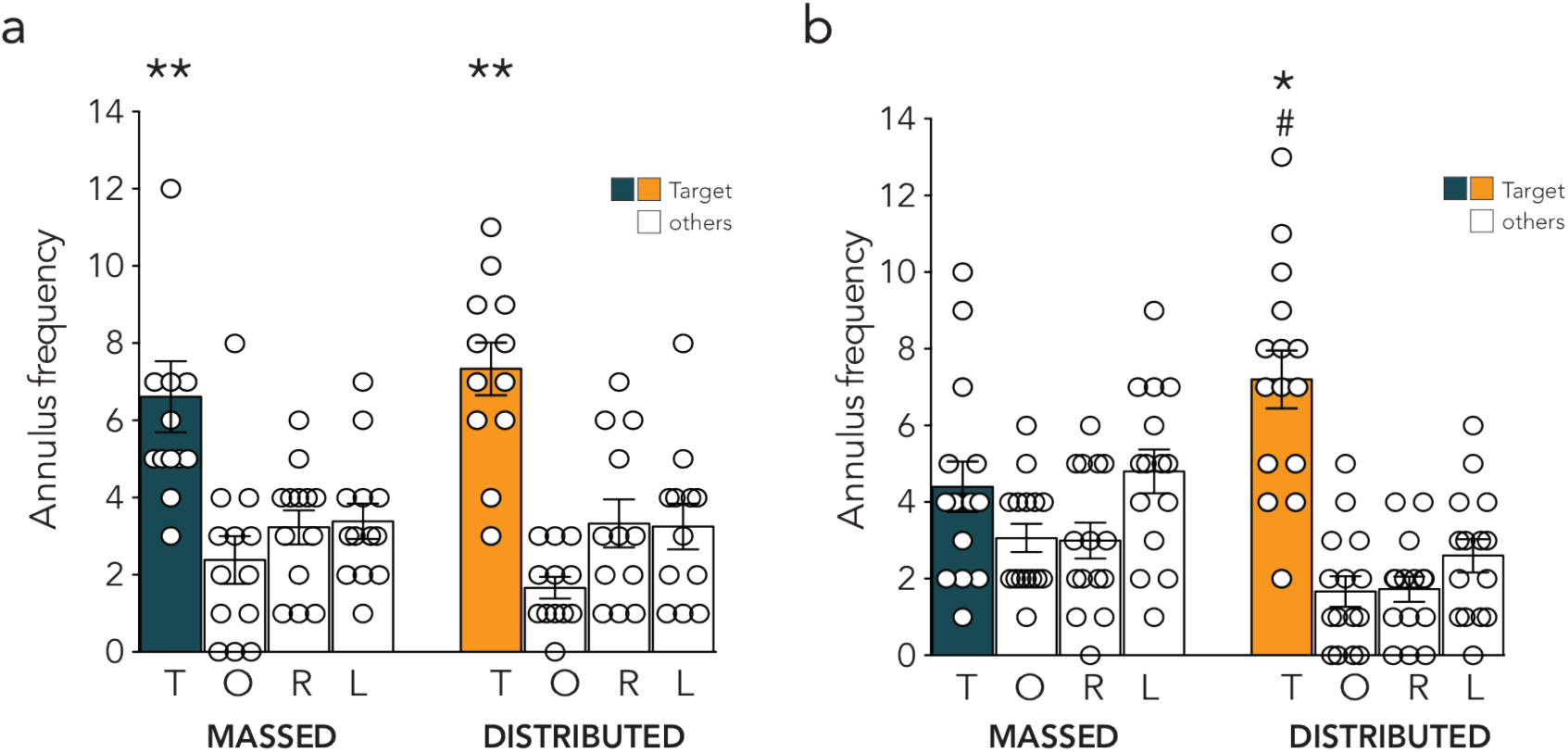
Distributed training increases memory stability. Mean annulus frequency ± SEM on test trial 24h **(a)** or 14 days **(b)** after massed or distributed training in the Morris water maze (24 h: twoway ANOVA repeated measure: quadrant F_(3,69)_=22.08, p< 0.0001; protocol F_(1,23)_= 0.00054, p = 0.9816; quadrant x protocol F_(3,69)_ = 0.4273, p = 0.7341; 14 days: two-way ANOVA repeated measure: quadrant F_(3,84)_ = 17.32, p < 0.0001; protocol F_(1,28)_ = 3.35, p = 0.077; quadrant x protocol F_(3,84)_= 8.335, p < 0.0001). *p<0.05 target vs right, opposite, left (within group, Tukey HSD); # p<0.05 target vs target (between group, Tukey HSD).

**Extended Data Fig. 2.**
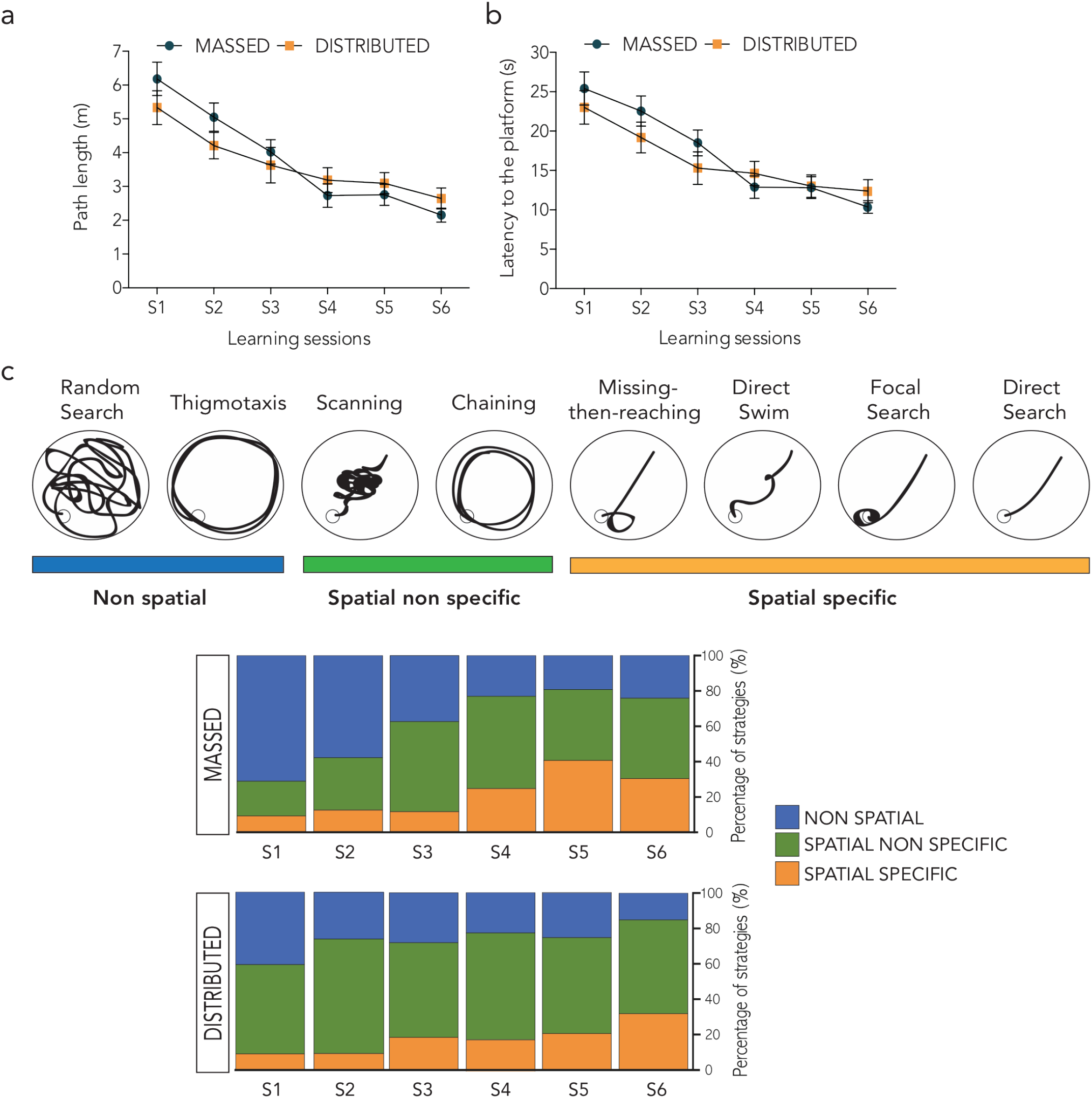
Training and strategy deployment of mice trained with the massed or the distributed-training protocol. **a,** Path length expressed as mean distance ± SEM during spatial training in the Morris water maze of mice trained with the massed (n=28) or distributed protocol (n=27) (two-way ANOVA repeated measure: session F _(5, 265)_ = 24.38, p < 0.0001; protocol F_(1, 53)_ = 0.20 p = 0.6541; session x protocol F _(5, 265)_ = 1.57 p= 0.1688). **b,** Mean latency ± SEM to reach the platform of mice trained in the Morris water maze with the two protocols (two-way ANOVA repeated measure: session F _(5, 265)_ = 21.12 p < 0.0001; protocol F _(1, 53)_ = 0.39 p = 0.5355; session x protocol F _(5, 265)_ = 1.27 p = 0.2772). **c,** Schematic representation of the search strategies classified and color coded for each category. The graph shows the average prevalence of each category across sessions of mice trained with the two protocols. Comparison between the two groups did not reveal any significant difference in strategy deployment (multinomial regression).

**Extended Data Fig. 3.**
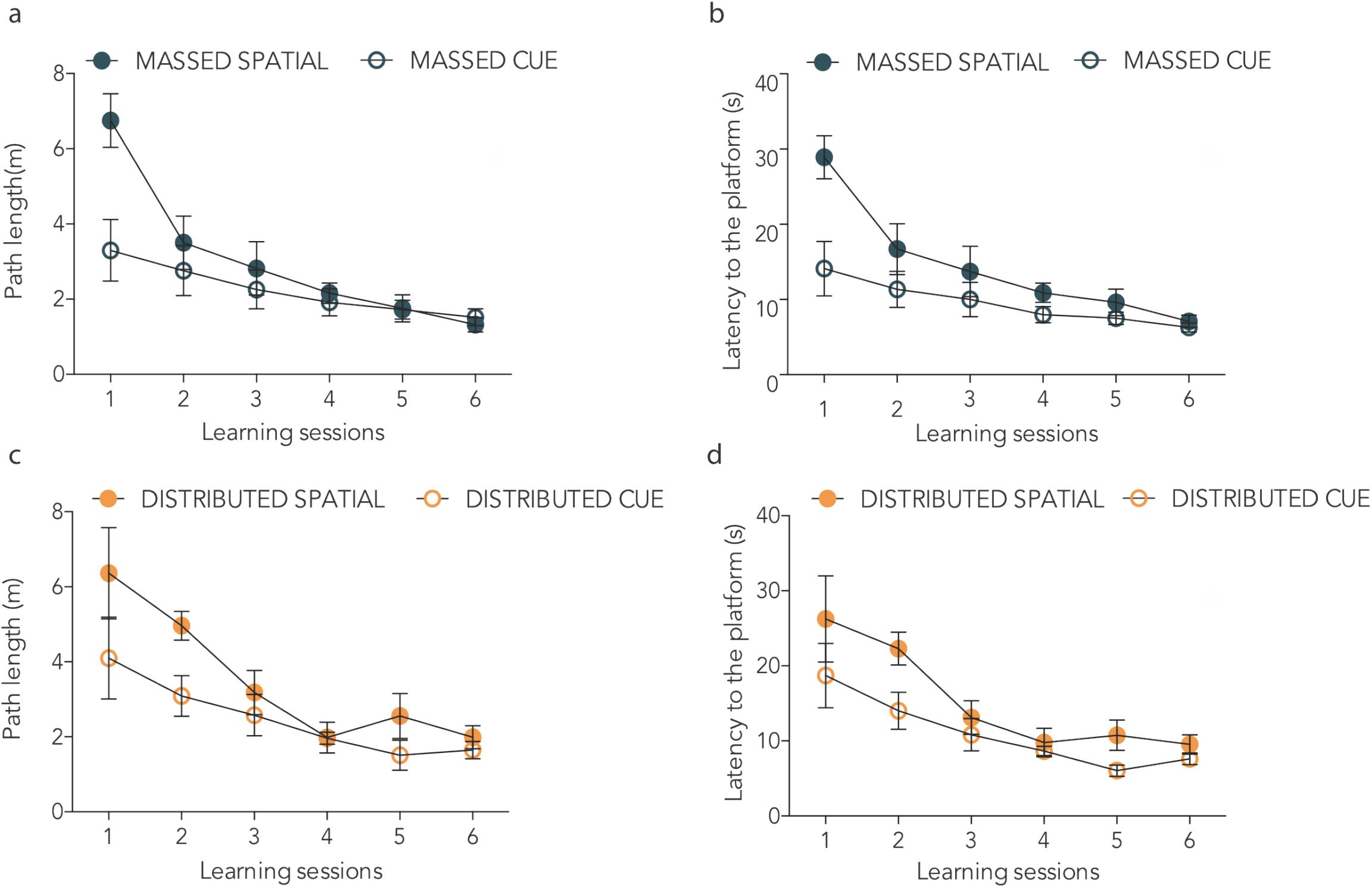
Learning curves for mice trained in the spatial or the cue version of the MWM with massed or distributed training protocols. **a,** Mean path length (m) ± SEM of mice massed trained in the sMWM or the cMWM (two-way ANOVA repeated measure: session F_(5,65)_=14.07, p<0.0001; protocol F_(1,13)_=4.02, p=0.0664; session x protocol F_(5,65)_=3.67, p=0.0055). **b,** Mean latency to reach the platform (s) ± SEM of mice massed trained in the spatial or cue version of the task (two-way ANOVA repeated measure: session F_(5,65)_=12.97, p<0.0001; protocol F_(1,13)_=6.97, p=0.0204; session x protocol F_(5,65)_=2.95, p=0.0184)**. c,** Mean path length (m) ± SEM of mice trained with the distributed protocol in the spatial or the cue version of the task (two-way ANOVA: session F_(5,70)_=10.23, p < 0.0001; protocol F_(1,14)_=6.7, p=0.0213; session x protocol F_(5,70)_=1.053, p=0.394). **d,** Mean latency to reach the platform (s) ± SEM of mice trained with the distributed protocol in the in the sMWM or the cMWM (two-way ANOVA repeated measure: session F_(5,70)_=10.77, p<0.0001; protocol F_(1,14)_=6.15, p=0.0265; session x protocol F_(5,70)_=0.7254, p=0.6067).

**Extended Data Fig. 4.**
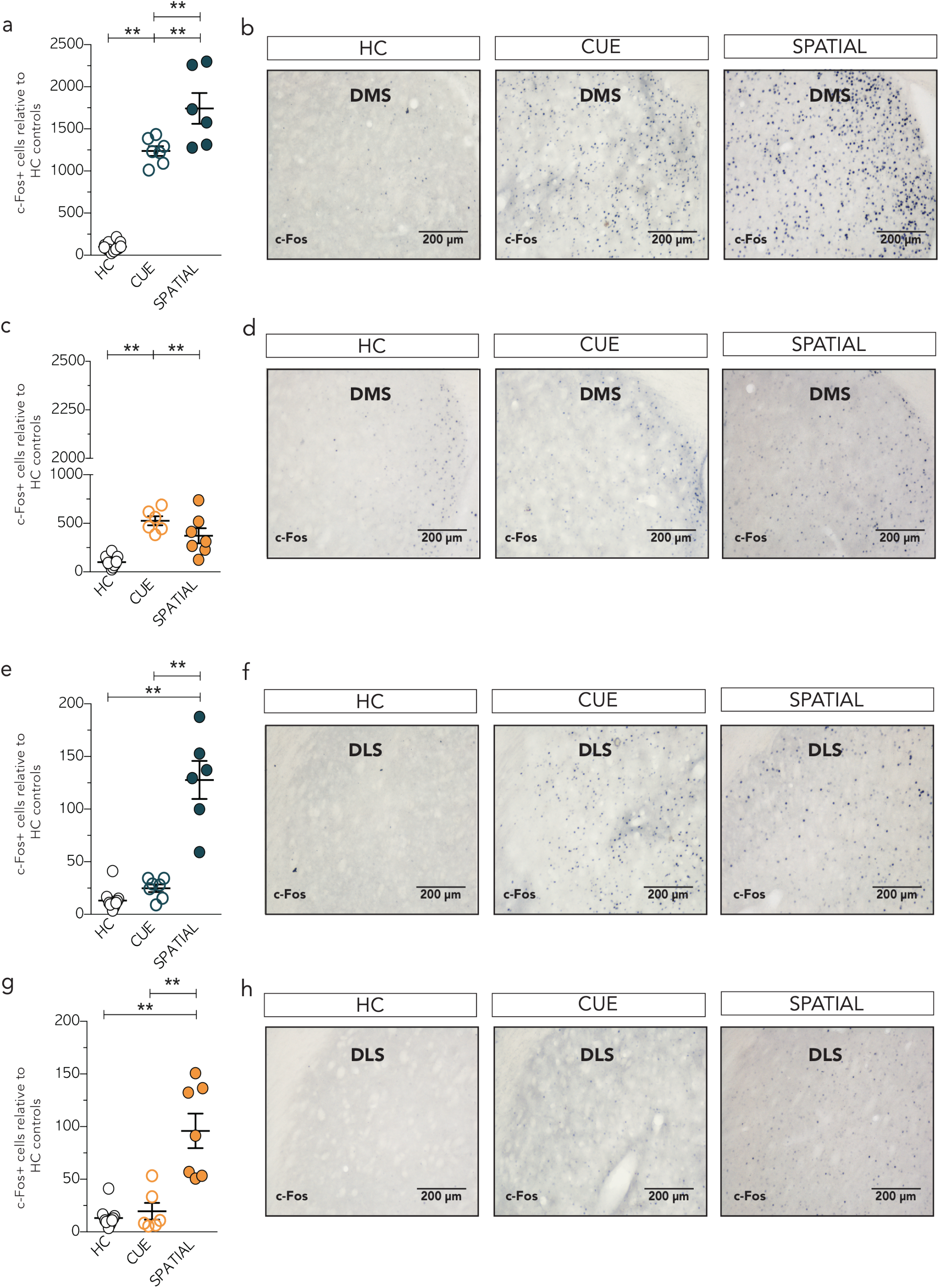
Spatial training induced c-Fos expression in the DMS and DLS after massed or distributed training. **a,** c-Fos labeling in the DMS was significantly higher in mice massed trained in the sMWM (n=6) compared to cMWM (n=7) or HC controls (n=12) (one-way ANOVA F_(2,22)_ = 117.8, p < 0.0001). **b,** Representative images of c-Fos immunoreactivity in the DMS in the three experimental groups. **c,** Distributed training significantly increased c-Fos expression in the DMS in both the cue (n=6) and the spatial (n=7) trained mice compared to the HC controls (one-way ANOVA F_(2,22)_ = 25.36, p< 0.0001). **d,** Representative images of c-Fos immunoreactivity in the DMS in the three experimental groups. **e**, Massed protocol induces a significantly higher c-Fos expression in the DLS following the sMWM compared to cMWM or HC controls (one-way ANOVA F_(2,22)_ = 54.39, p < 0.0001). **f**, Representative images of c-Fos immunoreactivity in the DLS in the three experimental groups. **g,** Distributed training in the sMWM induces a significantly higher c-Fos expression in the DLS compared to training in the cMWM or in HC controls (oneway ANOVA F_(2,22)_ =25.33, p < 0.0001). **h**, Representative images of c-Fos immunoreactivity in the DLS in the three experimental groups. **p<0.01 (Tukey HSD). Scale bars: 200 μm.

**Extended Data Fig. 5.**
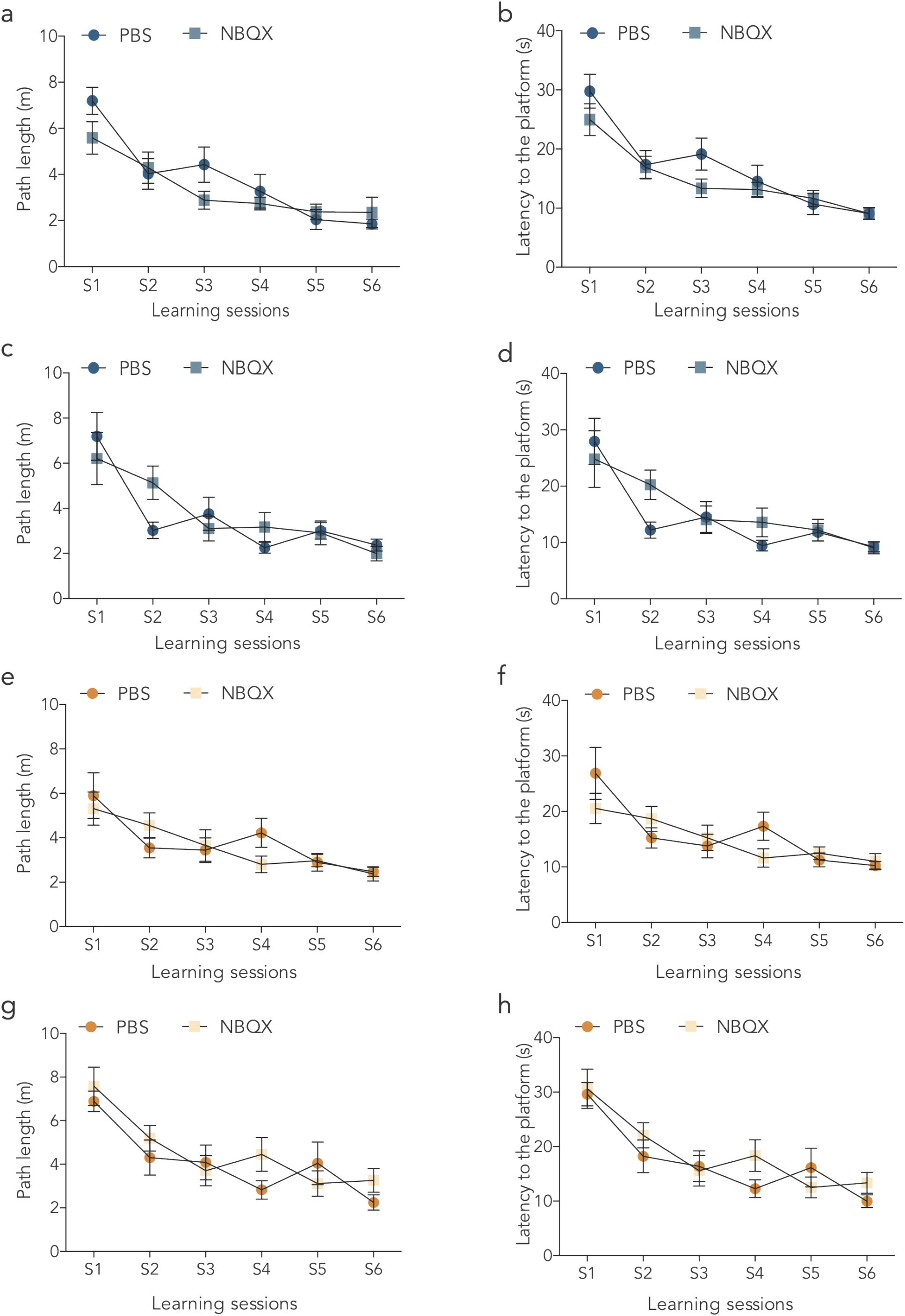
Training performance for massed- and distributed-training groups in the pharmacological experiments. **a,** Mean of path length (m) ± SEM after massed trained in the sMWM, before bilateral administrations in the DMS with PBS or NBQX (two-way repeated measure ANOVA of session F_(5,165)_=8.28, p<0.0001; treatment F_(1,33)_=1.050, p0.313; session x treatment F_(5,16)_ =1.659, p=0.1474). **b,** Mean of latency to reach the platform (s) ± SEM of mice massed trained in the sMWM before bilateral administrations in the DMS of vehicle or NBQX (two-way repeated measure ANOVA: session F_(5,165)_=24.56, p<0.0001; treatment F_(1,33_=1.406, p 0.2442; session x treatment F_(5,165)_=1.13, p=0.3487)**. c,** Mean of path length (m) ± SEM of massed trained mice in the sMWM before bilateral vehicle or NBQX administrations in the DLS (two-way repeated measure ANOVA: session F_(5,95)_=14.79, p<0.0001; treatment F_(1,19)_=0.103, p=0.7516; session x treatment F_(5,95)_=1.9, p=0.1009). **d,** Mean of latency to reach the platform (s) ± SEM of mice massed trained in the sMWM before bilateral DLS vehicle or NBQX administrations (two-way repeated measure ANOVA: session F_(5,95)_=14.26, p<0.0001; treatment F_(1,19)_=0.48, p=0.4952; session x treatment F_(5,95)_=1.5, p=0.1942). **e,** Mean of path length (m) ± SEM of mice trained with the distributed protocol in the sMWM before bilateral vehicle or NBQX administrations in the DMS (two-way repeated measure ANOVA: session F_(5,115)_ = 7.93, p < 0.0001; treatment F_(1,23)_ = 0.13, p = 0.7234; session x treatment F_(5,115)_ = 1.09, p = 0.3697). **f,** Mean of latency to reach the platform (s) ± SEM in the sMWM of the distributed trained mice before bilateral vehicle or NBQX administrations in the DMS (two-way repeated measure ANOVA: session F_(5,115)_=9.22, p<0.0001; treatment F_(1,23)_=0.33, p=0.5706; session x treatment F_(5,115)_=1.804, p=0.1174). **g,** Mean of path length (m) ± SEM of mice trained with the distributed protocol in the sMWM before bilateral vehicle or NBQX administration in the DLS (two-way repeated measure ANOVA: session F_(5,135)_=12.36, p<0.0001; treatment F_(1,27)_=0.87, p=0.3591; session x treatment F_(5,135)_=1.14, p=0.34). **h,** Mean of latency to reach the platform (s) ± SEM of mice trained on the distributed protocol in the sMWM before bilateral vehicle or NBQX administrations in the DLS (two-way repeated measure ANOVA: session F_(5,135)_=15.61, p<0.0001; treatment F_(1,27)_=0.65, p=0.4282; session x treatment F_(5,135)_=1.12, p=0.3545).

**Extended Data Fig. 6.**
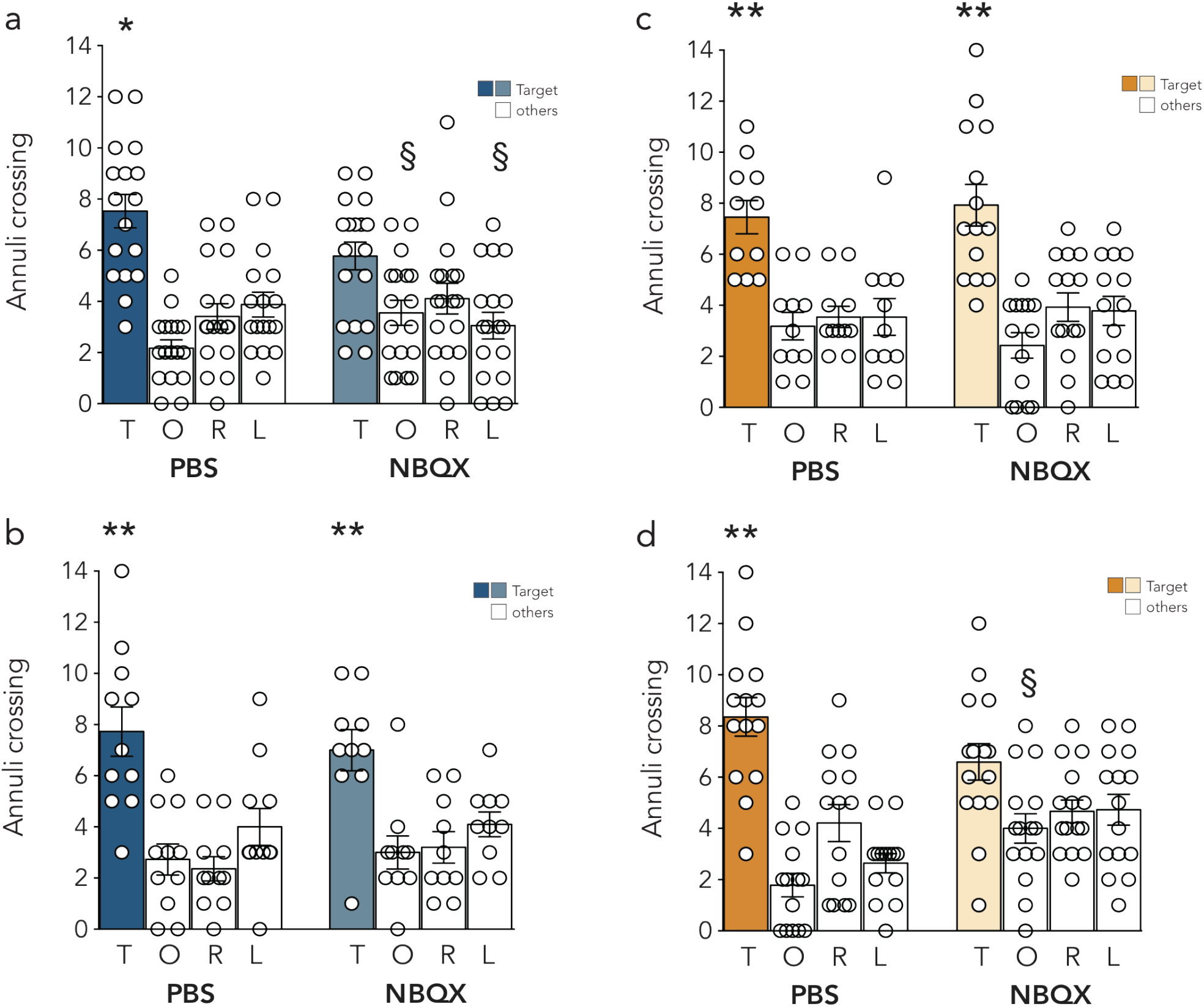
Pre-test bilateral inhibition of the DMS or the DLS following massed or distributed training in the sMWM. **a,** Mean of annulus frequency ± SEM on probe trial 24h after massed training of mice administered pre-test in the DMS with vehicle or NBQX (two-way ANOVA repeated measure: annuli F_(3,99)_=21.76, p<0.0001; treatment F_(1,33)_=0.099, p=0.7553; annuli x treatment F_(3,99)_=3.88, p=0.0114)**. b,** Mean of annulus frequency ± SEM on probe trial test, 24h after massed training, of mice administered with either vehicle or NBQX in the DLS (two-way ANOVA repeated measure: annuli F_(3,57)_=16.59, p<0.0001; treatment F_(1,19)_=0.13, p=0.72; annuli x treatment F_(3,57)_=0.3767, p=0.77). **c,** Mean of annulus frequency ± SEM on probe trial of distributed trained mice after bilateral DMS vehicle or NBQX administrations (two-way ANOVA repeated measure: annuli F_(3,69)_=21.26, p<0.0001; treatment F_(1,23)_=0.078, p=0.7815; annuli x treatment F_(3,69)_=0.3585, p=0.7832). **d,** Mean of annulus frequency ± SEM on probe trial 24h after last training, on the distributed protocol, of mice administered pre-test vehicle or NBQX in the DLS (two-way ANOVA repeated measure: annuli F_(3,81_=21.21, p<0.0001; treatment F_(1,27)_=4.13, p=0.0521; annuli x treatment F_(3,81)_=4.54, p=0.0054). *p<0.05 target vs right, opposite, left (within group, Tukey HSD); § p<0.05 *vs* target (within groups, Tukey HSD).

**Extended Data Fig. 7.**
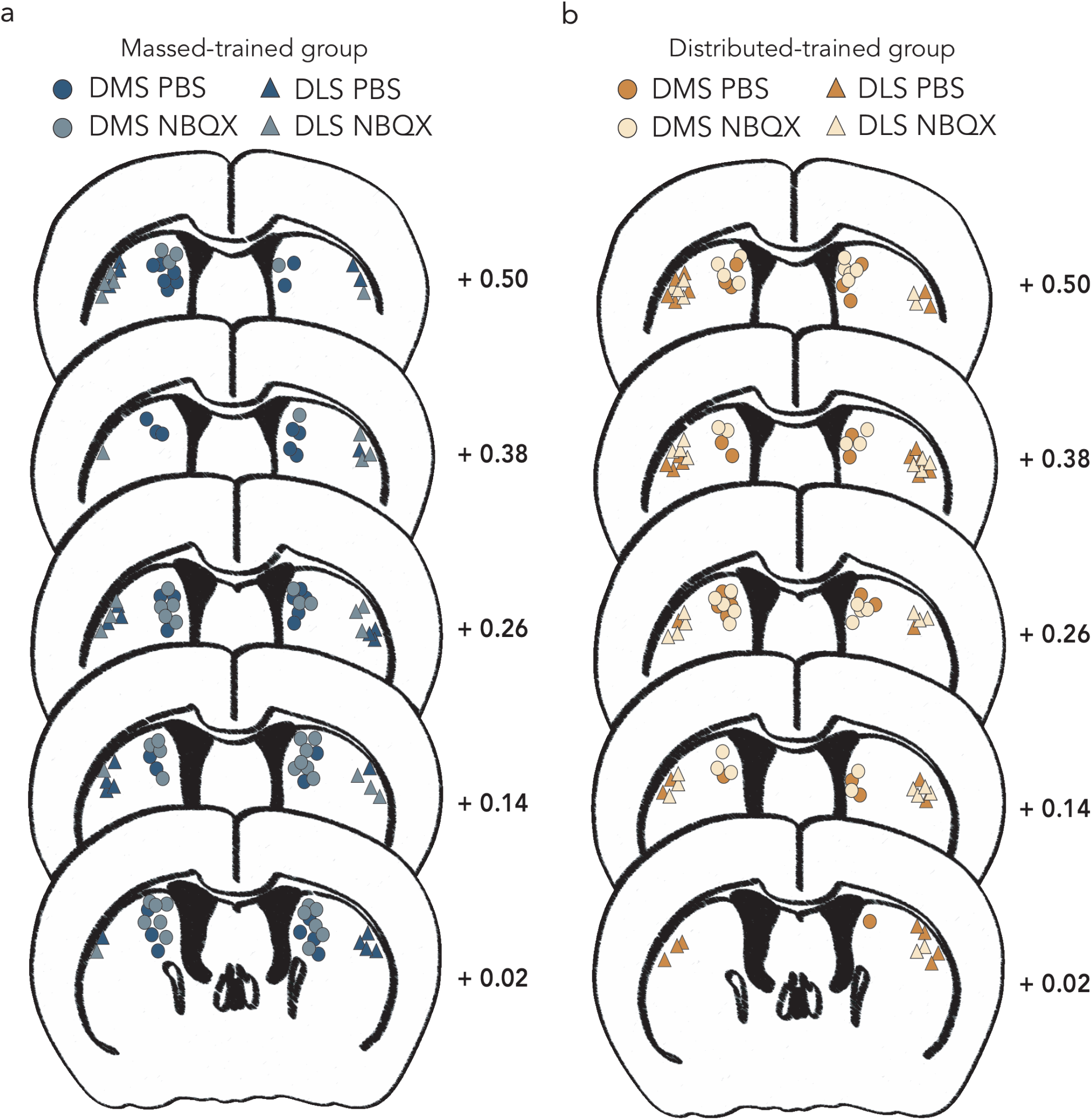
Schematic drawings of the injection sites for the pharmacological experiments. Each symbol represents the site of injection for one animal. **a**, Massed training groups. Pre-test DMS vehicle (blue circles, n = 17) and NBQX (grey circles, n = 18); pre-test DLS vehicle (blue triangles, n = 11) and NBQX (grey triangles, n = 10). **b**, Distributed training groups. Pre-test DMS vehicle (orange circles, n = 11) and NBQX (yellow circles, n = 14); pre-test DLS vehicle (orange triangles, n = 14) and NBQX (yellow triangles, n = 15). Coordinates are expressed as mm from bregma.

**Extended Data Fig. 8.**
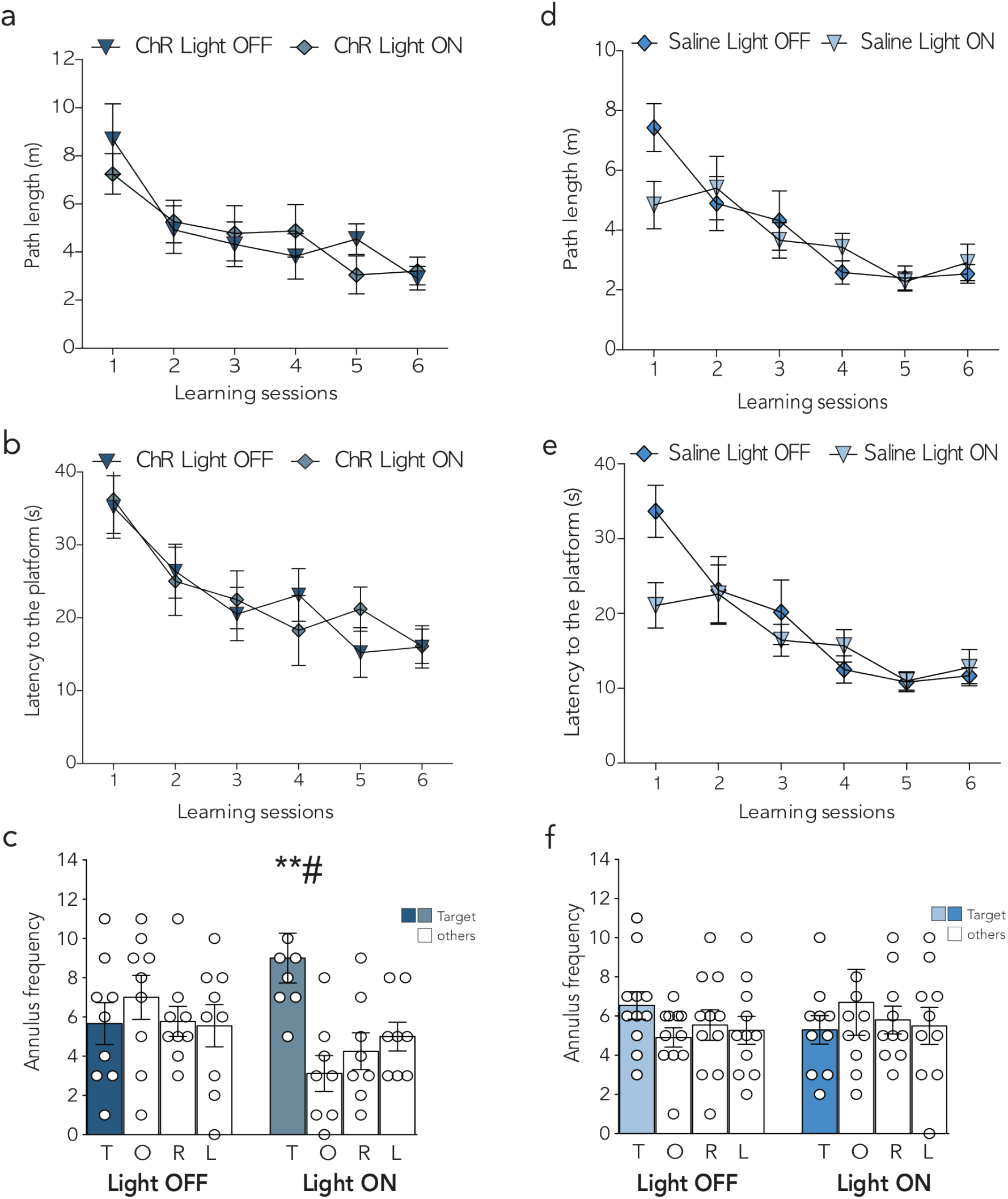
Light delivery in the DLS improved memory performance 14 days after massed training only in ChR2(C128S/D156A) expressing mice. **a,** Mean path length (m) ± SEM during massed training of ChR-eYFP expressing mice with (Light ON) or without (Light OFF) light delivery (two-way ANOVA: session F_(5,75)_ = 7.929, p < 0.0001; treatment F_(1,15)_ = 0.0286, p = 0.8679; session x treatment F_(5,75)_ = 0.7919, p = 0.5589). **b,** Mean of latency (s) ± SEM during massed training of stimulated (Light ON) and unstimulated (Light OFF) ChR-eYFP expressing mice (two-way ANOVA repeated measure: session F_(5,75)_ = 8.054, p < 0.0001; treatment F_(1,15)_ = 0.227, p = 0.64; session x treatment F_(5,75)_ = 0.3382, p = 0.8883). **c**, Mean of annulus frequency ± SEM on probe trial of stimulated and unstimulated ChR-eYFP expressing mice (twoway ANOVA: annuli F_(3,45)_ = 2.513, p = 0.0705; treatment F_(1,15)_ = 0.7932, p = 0.3872; annuli x treatment F_(3,45)_ = 4.568, p = 0.0071). **d,** Mean path length (m) ± SEM during massed training of saline injected groups with (Light ON) or without (Light OFF) light stimulation (two-way ANOVA repeated measure: session F_(5,95)_ = 10.66, p < 0.0001; treatment F_(1,19)_ = 0.3007, p = 0.5898; session x treatment F_(5,95)_ = 1.874, p = 0.1060). **e,** Mean of latency (s) ± SEM during massed training of stimulated and unstimulated saline-injected mice (twoway ANOVA repeated measure: session F_(5,95)_ = 11.78, p < 0.0001; treatment F_(1,19)_ = 0.8741, p = 0.361; session x treatment F_(5,95)_ = 2.226, p = 0.058). **f**, Mean of annulus frequency ± SEM on probe trial of stimulated and unstimulated saline injected groups (two-way ANOVA repeated measure: annuli F_(3,57)_ = 0.0.126, p = 0.9443; treatment F_(1,19)_ = 0.2011, p = 0.6589; annuli x treatment F_(3,57)_ = 0.91, p = 0.44). *p<0.05 target vs right, opposite, left (within group, Tukey HSD); # p<0.05 target vs target (between group, Tukey HSD).

**Extended Data Fig. 9.**
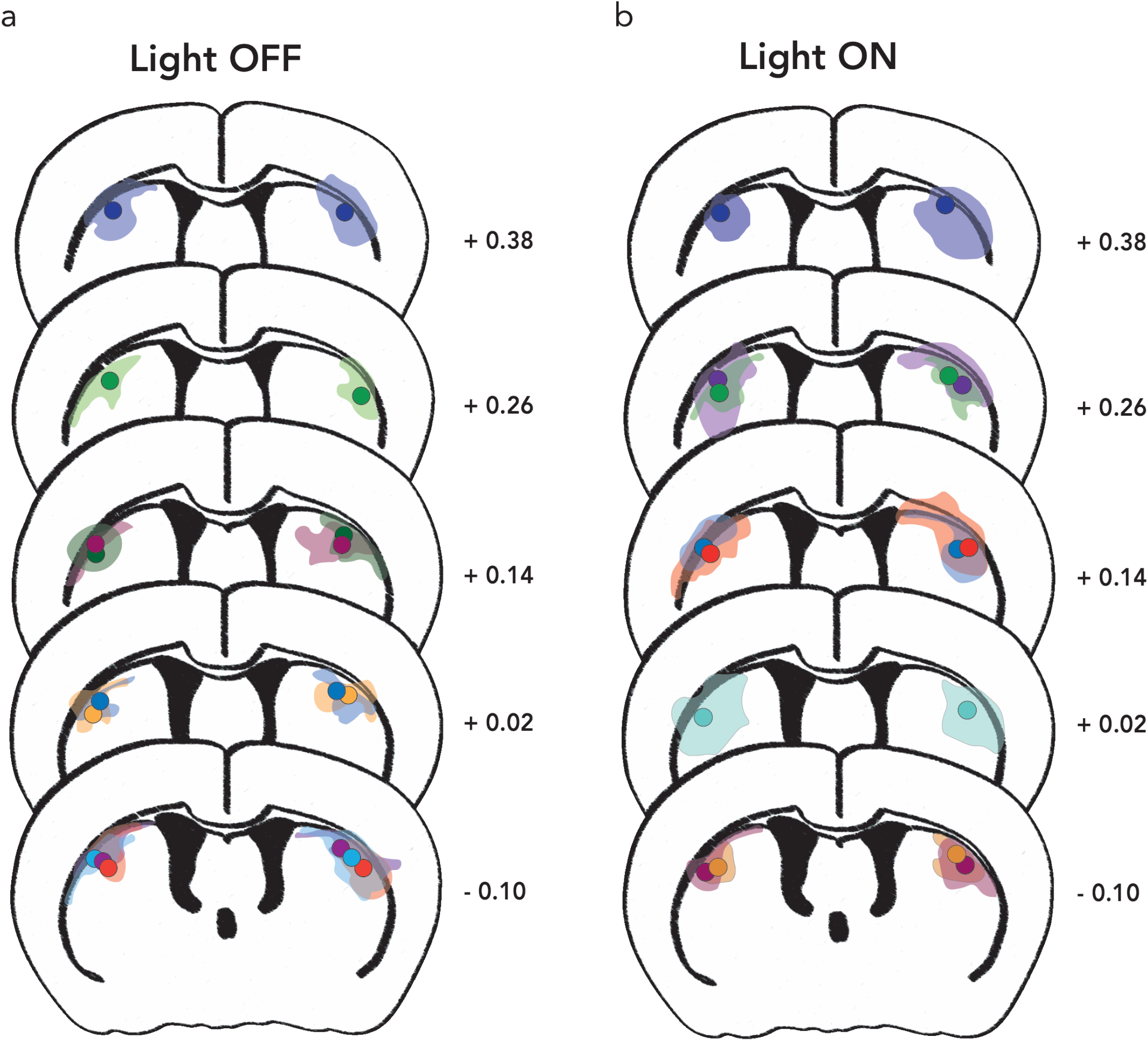
AAV:: ChR2(C128S/D156A) diffusion in the DLS of stimulated and unstimulated mice. Schematic representation of eYFP protein expression and fiber placements in **a,** the light-OFF (n=9) and **b,** the light-ON (n=8) groups. For each animal eYFP expression (lighter) and the fiber placement (darker dots) are represented in the same color. Coordinates are expressed as mm from bregma.

**Extended Data Fig. 10.**
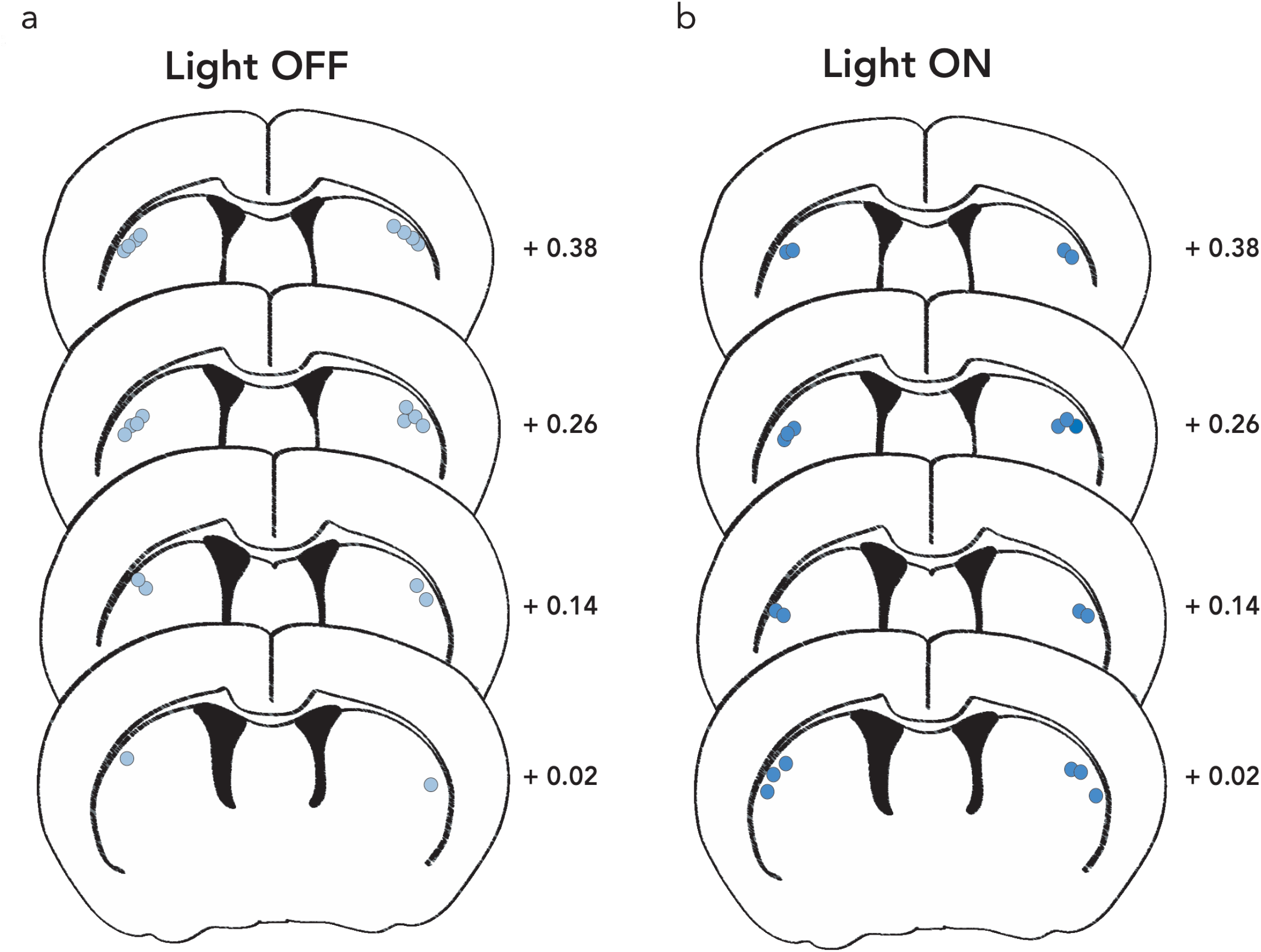
Schematic drawing of the fiber placements in stimulated and unstimulated saline controls. Schematic representation of the site of injection for saline-injected mice **a,** without light delivery (LIGHT OFF) or **b,** with light delivery (LIGHT ON). Coordinates are expressed as mm from bregma.

**TABLE S1.** Multinomial logistic regression analysis for navigational strategies across the learning sessions.

|  | B | Standard Error | Wald | gl | Sign. |
| --- | --- | --- | --- | --- | --- |
| Intercept | 0.253 | 0.211 | 1.44 | 1 | 0.229 |
| S1 * strategy | -0.154 | 0.102 | 2.28 | 1 | 0.131 |
| S2 * strategy | -0.147 | 0.104 | 2.00 | 1 | 0.157 |
| S3 * strategy | -0.148 | 0.112 | 1.73 | 1 | 0.189 |
| S4 * strategy | -0.110 | 0.115 | 0.91 | 1 | 0.340 |
| S5 * strategy | -0.067 | 0.121 | 0.31 | 1 | 0.578 |
| S6 * strategy | -0.179 | 0.126 | 2.01 | 1 | 0.156 |

